# UniLoc: A universal protein localization site predictor for eukaryotes and prokaryotes

**DOI:** 10.1101/252916

**Authors:** Hsin-Nan Lin, Ching-Tai Chen, Ting-Yi Sung, Wen-Lian Hsu

## Abstract

There is a growing gap between protein subcellular localization (PSL) data and protein sequence data, raising the need for computation methods to rapidly determine subcellular localizations for uncharacterized proteins. Currently, the most efficient computation method involves finding sequence-similar proteins (hereafter referred to as *similar proteins*) in the annotated database and transferring their annotations to the target protein. When a sequence-similarity search fails to find similar proteins, many PSL predictors adopt machine learning methods for the prediction of localization sites. We proposed a universal protein localization site predictor - UniLoc - to take advantage of implicit similarity among proteins through sequence analysis alone. The notion of related protein words is introduced to explore the localization site assignment of uncharacterized proteins. UniLoc is found to identify useful template proteins and produce reliable predictions when similar proteins were not available.

## INTRODUCTION

The subcellular localization of a protein provides crucial information regarding the protein’s biological function, and can be used to help to annotate newly sequenced genomes, to verify experimental results, and to identify potential drug targets [1]. Biologists world-wide have contributed localization data, but the gap between localization data and protein sequence data continues to increase, raising the need for computation methods to rapidly predict subcellular localization for uncharacterized proteins based on sequence information alone. The SwissProt database currently contains more than a half million manually annotated and reviewed protein records. The annotations are taken from the literature and curator-evaluated computational analysis, and computational analysis has played an important role in PSL annotation. Around 90% of protein records in Swiss-Prot database are annotated through sequence similarity. For protein sequences without obvious sequence-similar proteins, a PSL predictor helps to provide plausible information quickly.

The most efficient approach to predict the subcellular localization sites for a target protein is to use similar proteins in the Swiss-Prot database as templates, and to transfer their annotations to the target sequence. When a similarity search fails to find such proteins, various protein subcellular localization (PSL) predictors use different features including amino acid composition[2–8], sequence patterns[9–12], physical-chemical features of protein sequences[11, 13–18], biological features from the literature and databases[4, 19–23], Gene Ontology annotations[24–31], and protein-protein interactions[32, 33]. Some approaches develop algorithms to combine the prediction results of existing PSL predictors and produce the consensus prediction, such as SUBAcon [34].

Many PSL predictors were built specifically for a target species due to the difference in subcellular locations of similar or homologous proteins. Users choose a target species to avoid noisy sequences. However, we found that proteins with the same localization sites in different species may share an implicit similarity, and a PSL predictor could take advantage of such similarities among proteins regardless of their species. It is also observed that sophisticated machine learning algorithms in state-of-the-art PSL predictors can make it difficult to trace the prediction back the most important features leading to a particular prediction [31]. Such information may be helpful for biologists to further determine what features are likely responsible for the localization.

In this study, we present UniLoc for PSL prediction. UniLoc is a successor of KnowPred_site_ [9]. The major improvements of UniLoc include: 1) UniLoc is able to predict single‐ and multilocalized proteins, whereas KnowPred_site_ could only estimate a probability of each localization site. So, it is difficult for KnowPred_site_ to evaluate its performance on the multi-localized protein prediction and to compare with existing methods. The performance of UniLoc is compared with state-of-the-art methods and a PSIBLAST-based predictor. 2) UniLoc identifies template proteins based on implicit local similarities and makes the prediction with the identified significant templates, whereas KnowPred_site_ makes predictions with all possible proteins. 3) UniLoc is trained and evaluated with non-redundant proteins of eukaryotes and prokaryotes, whereas KnowPred_site_ was trained and evaluated with a set of redundant eukaryotic proteins. UniLoc was trained with a non-redundant data set consisting of 14754 eukaryotic proteins and 2585 prokaryotic proteins covering 1377 species. It not only predicts the localization sites for a protein sequence, but also provides potentially related proteins. In addition, UniLoc offers confidence scores to indicate the reliability of the prediction. The UniLoc web server is available at http://bioapp.iis.sinica.edu.tw/UniLoc/.

## MATERIALS AND METHODS

### Data sets

A set of 538,010 proteins were obtained from UniProt/SwissProt database. Proteins with fewer than 50 amino acids or without experimentally determined PSL annotations were removed, leaving a subset of 65,506 proteins. Localization annotations were further categorized according to the taxonomy described in [35]. To construct a non-redundant benchmark data set, PSI-CD-HIT was used to exclude proteins that shared greater than 30% sequence identity with any other protein. Subsequently, localization sites were not considered if the number of proteins in a category was less than 30. Eventually, we have two benchmark data sets, one containing 14,754 eukaryotic proteins (denoted as *SP-Euk*), and the other containing 2,585 prokaryotic proteins (denoted as *SP-Prok*).

The numbers of localization sites of *SP-Euk* and *SP-Prok* are 16 and 5, respectively. For *SP-Euk*, 8798 proteins (~60%) are single-localized and 5956 proteins (~40%) are multi-localized. In contrast, most prokaryotic proteins (~92%) in *SP-Prok* are single-localized. *SP-Euk* and *SP-Prok* respectively embrace 836 and 541 species.

### Similar words vs. related words

In the field of computational linguistics, n-gram models have been shown to achieve good results for document classification [36, 37]. Similar techniques have also been applied to classify and characterize protein sequences. N-grams in a protein sequence are a set of substrings of a fixed length *n* and are often referred to as protein words. Thus a protein sequence can be viewed as a text written in the protein language consisting of 20 amino acids. The mapping of protein words to protein functions is conceptually similar to the mapping of words to meaning.

BLAST’s search algorithm [38] uses a sliding window to generate all possible words (referred to as the *initial word set*) that appear throughout the target sequence. Each word is then used as a seed to identify potentially similar proteins. To increase the sensitivity of protein words, BLAST defines similar words with respect to a word in the initial word set based on the alignment score of two words.

Combining the idea of similar words and the mapping of protein words to protein functions, we present the concept of related words to extend similar words. Given a protein sequence *p*, we perform a PSI-BLAST search against NCBI NR database. PSI-BLAST generally returns a number of sequence alignments (HSP with e-value < 0.001) between *p* and its similar proteins *sp*. All the words in *p* and *sp* are extracted by a sliding window of size n. Given a word *w* (without gaps) in *p*, its *related word* is defined as the word *w’* (without gaps) in *sp* that is aligned with *w*. It is noteworthy that the same word *w* can be related to a number of words *w’* if it is aligned with different protein sequences. Unlike similar words which were defined based on the alignment score of the word pair alone, related words are defined based on a significant sequence alignment of two proteins. In the example of Figure 1, EWQLVLH and DFDMVL are related words, and so are HVWAKVE and KCWGPVE. We use a sliding window of size *n* to screen the alignment to extract all related word pairs. No gap is allowed in a related word pair, thus every word represents an *n*gram. In the example of Figure 1, we can collect 34 related word pairs of length 7 from the alignment.

**Figure 1.**
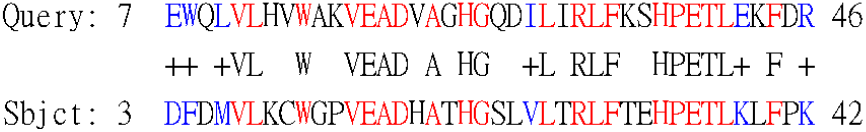
An example of significant sequence alignment. The alignment shows that the sequence fragment from position 7 to position 46 of the query sequence is highly similar with that from position 3 to position 42 of the subject sequence.

The choice of word length *n* is a trade-off between specificity and sensitivity, i.e., a larger *n* yields high specificity, whereas a smaller *n* produces high sensitivity. In sequence similarity identification, a smaller *n* is preferred. The word length in this study was empirically set as 7 to achieve balance between specificity and sensitivity [9, 11, 39].

Given a protein sequence *p*, we apply PSI-BLAST[40] to search against NCBI nr database and obtain a group of high-scoring segment pairs (HSPs). Each HSP represents a pairwise sequence alignment between *p* and its similar protein *sp*. Every word *w* in *p* may obtain a number of distinct related words *w’* in *sp*. If the localization sites of protein *p* have been experimentally determined, the information can be hypothetically transferred to each of its related words. In this way, we can build a word dictionary, called *RelatedDict* given a group of proteins with subcellular localization information and their HSPs. An example of a related word entry in *RelatedDict* is shown in Table 1, where protein sources represent those proteins which contain an initial word related to the word entry (MSQTAPS) and has been annotated with localization information. In this example, there are four protein sources, three of which are single-localized into the cytoplasm, and one is multilocalized into the cytoplasm and nucleus.

**Table 1.**
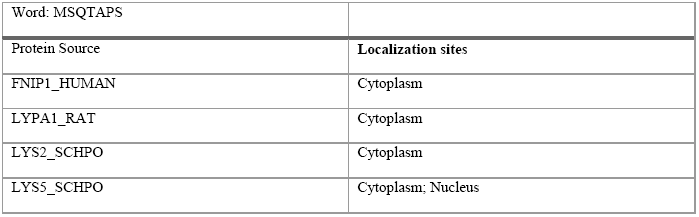
An example of a related word entry in RelatedDict. The word, MSQTAPS, is related to the words of the four protein sequences and it inherits their location information.

### Identification of template proteins using related words and localization prediction

Given a target protein *p* and its related word set extracted from all the HSPs, each related word is used to match against the *RelatedDict*. If a word *w* matches a dictionary entry exactly, it means every protein source in this word entry shares a related word with *p*. UniLoc identifies template proteins based on the number of shared related words. Suppose there are *N* related words in the word set of *p* matched against *RelatedDict* and a protein *q* shares *f* related words with *p*, we define the *intersection percentage* of *q* over *p* as (100 × *f* / N)%. Protein *q* is considered to be a *template protein* if its intersection percentage is over 10%. In case a template protein is found, UniLoc predicts the localization sites of *p* based on the top 5 template proteins. Assume *t_1_, t_2_, …, t_n_* (*n* ≤ 5) are the top *n* template proteins, the confidence score *CS*(*s_i_*) for localization site *s_i_* is defined as the ratio of its frequency to *n*. For example, if three out of five template proteins are localized into the nucleus, then *CS*(*nucleus*) is 3/5 = 60%.

If none of template proteins can be found, UniLoc predicts the localization sites of *p* based on the individually matched related words. Given a matched word *w*, the confidence score *CSw*(*s_i_*) for localization site *s_i_* is defined as the ratio of its frequency to the number of protein sources of the entry in *RelatedDict.* Take the related word in Table 1, as an example, *CSw*(*cytoplasm*) is 100 (= 100 × 4 / 4), whereas the *CSw*(*nucleus*) is 25 (=100 × 1 / 4). If there are *N* matched word entries, the confidence score *CS*(*s_i_*) is the summation of all *CSw*(*s_i_*) divided by *N*, so that *CS*(*s_i_*) is normalized between 0 and 100.

The prediction result is derived from the confidence scores of all localization sites. For a single-localized protein, the prediction is simply the localization site with the highest confidence score; however, for a multi-localized protein, determining the prediction is more difficult because it is unclear how many localization sites should be reported. To handle this situation, we modify the multi-localized confidence score described in [11], denoted as MLCS(*s_i_*), to determine the prediction according to the confidence scores of all the sites. The MLCS(*s_i_*) for localization site *s_i_* is defined as follows:

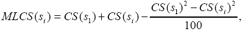

where *s*_1_ is the localization site with the highest score. UniLoc always reports *s*_1_ (i.e., the top1) in the prediction and reports the remaining localization site *s_i_* if the following two criteria are satisfied:

1) CS(*s_i_*) > *CSthreshold* and 2) MLCS(*s_i_*) > *MLCSthreshold*. We will discuss the performance of different *CSthreshold* and *MLCSthreshold* in the results section.

### A PSI-BLAST based prediction method

Since PSI-BLAST is a powerful tool for sequence similarity search, it has been widely used in the identification of protein templates for function and structure prediction. The most significant feature of PSI-BLAST is that it iteratively searches a protein database for similar sequences and uses position-specific scoring matrices to identify distantly related sequences. PSI-BLAST is more sensitive than BLAST, therefore we implemented a predictor based on PSI-BLAST search results. Given a target protein *p*, we perform a PSI-BLAST search with iteration 3 against the benchmark data set (*SP-Euk* or *SP-Prok*) and use the best five proteins as templates for localization assignment. The assignment strategy is similar to that of UniLoc, except that UniLoc identifies templates by looking up related words in the *RelatedDict.* If PSI-BLAST fails to find any protein, no prediction will be made. The performance of this PSI-BLAST based predictor is used as the baseline for the benchmark data sets.

### Evaluation metrics

In this study, we performed 10-fold cross-validation on *SP-Euk* and *SP-Prok* to estimate the performance of UniLoc. In each experiment, a *RelatedDict* was compiled using one of the two data sets and PSL annotations are excluded if they belong to the proteins in the test set. To evaluate the prediction result, the *Top1 success rate*, *precision*, *recall*, and *F-score* were calculated. Top1 success rate is defined as the precision of the localization site with the highest confidence score in predictions; precision is defined as TP / (TP + FP); recall as TP / (TP + FN), where TP, FP, and FN are the numbers of true positives (correctly predicted localization sites), false positives (incorrectly predicted localization sites), and false negatives (missing localization sites), respectively. And F-score is defined as (2 × precision × recall) / (precision + recall).

## RESULTS

### Efficacy of protein related words

Protein related words are the basic units we used for localization site prediction. To demonstrate the efficacy of related words, we analyse the average number of common related words between two single-localized proteins. In this analysis, we randomly selected 10,000 protein pairs with the same localization site and another 10,000 protein pairs with different localization sites.Figure 2 shows the average number of common related words between these protein pairs (i.e., same localization site vs. different localization site). It is clear that protein pairs with the same localization share more related words than those with different localization. For example, the average number of common related words between two Golgi apparatus localized proteins is 1486; however, the average number of common related words between two proteins, one localized in Golgi apparatus and the other localized somewhere else, is only 72. This suggests that Golgi apparatus localized protein sequences are more conserved. The average number of common related words between two extracell localized proteins is 59; the average number of common related words between two proteins, one localized into extracell and the other localized somewhere else, is 19. This implies that extracell localized protein sequences are more diverse.

**Figure 2.**
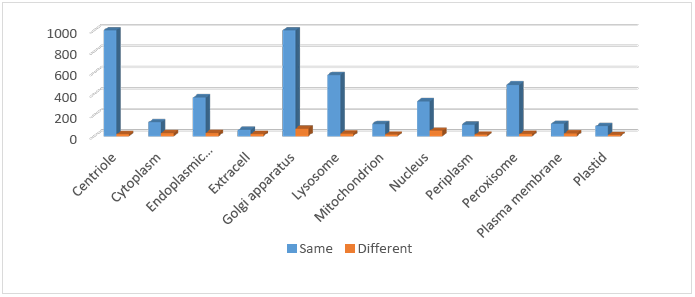
The average number of common related words analysis on protein pairs with the same or different localization site.

### CSthreshold and MLCSthreshold and overall performance on benchmark data sets

We use the two thresholds, CSthreshold and MLCSthreshold to determine what localization sites should be outputted. To identify a proper combination of the two parameters and avoid overfitting the benchmark data sets, we first evaluated the F-score of different CSthreshold values with a fixed MLCSthreshold of 0 to identify the best CSthreshold, and then evaluated the F-score of different MLCSthreshold values with the best CSthreshold. Table 2, summarizes the performance evaluation on *SP-Euk*. It shows that a CSthreshold of 30 and an MLCSthreshold of 60 produce the best performance. We use the same thresholds to estimate the performance of UniLoc on *SP-Prok*. The detailed performance of UniLoc and the PSI-BLAST-based predictor on the two benchmark data sets is shown in Table 3. UniLoc significantly outperforms the PSI-BLAST-based predictor, suggesting that related words can help detect implicit similarities and identify more template proteins with the same localization sites.

**Table 2.** Performance of UniLoc with different combinations of CSthreshold and MLCSthreshold

| CStreshold + MLCStreshold (0) | 10 | 20 | 30 | 40 | 50 | 60 | 70 | 80 | 90 |
| --- | --- | --- | --- | --- | --- | --- | --- | --- | --- |
| **F-score** | 68.3 | 67.5 | **68.6** | 67.8 | 67.4 | 66.9 | 66.5 | 66.4 | 66.3 |
| MLCStreshold + CStreshold (30) | 10 | 20 | 30 | 40 | 50 | 60 | 70 | 80 | 90 |
| **F-score** | 68.6 | 68.6 | 68.6 | 68.6 | 68.8 | **68.9** | 68.7 | 67.9 | 67.3 |

**Table 3.** The overall performance on the benchmark data sets

|  | *SP-Euk* |  | *SP-Prok* |  |
| --- | --- | --- | --- | --- |
|  | UniLoc | PSI-BLAST<br>predictor | UniLoc | PSI-BLAST<br>predictor |
| **Top1 success rate** | **71.9** | 57.7 | **66.2** | 50.7 |
| **Precision** | **65.0** | 51.4 | **71.2** | 49.7 |
| **Recall** | **73.2** | 58.1 | **81.5** | 51.8 |
| **F-score** | **68.9** | 54.5 | **76.0** | 50.7 |

### Similarity among proteins from different species

We assume proteins with the same localization sites may share implicit similarity. To verify this hypothesis, we estimated UniLoc’s performance with the constraint that prediction is made only using the annotation from proteins within the same species and compared to the performance without this constraint. From Table 4, it is clear that UniLoc’s performance is much better without the species constraints. Thus we can expect that the more proteins involved in the related word dictionary, the better UniLoc can predict for a target protein.

**Table 4.** Performance comparison on *SP-Euk* with/without the species constraints.

| *SP-Euk* | *No species constraints* | *With species constraints* | *Difference* |
| --- | --- | --- | --- |
| **Top1 success rate** | **71.9** | 59.7 | +12.2 |
| **Precision** | **65.0** | 54.5 | +10.5 |
| **Recall** | **73.2** | 64.7 | +8.5 |
| **F-score** | **68.9** | 59.2 | +9.7 |

### The reliability of confidence scores

Confidence scores play an important role in computational analysis and are indicators of prediction reliability. To estimate the efficacy of UniLoc’s confidence scores, we analyse the relationship between confidence scores and the corresponding precisions. From Figure 3, it can be observed that there is a strong positive correlation between the two factors. The correlation coefficient is 0.95 suggesting that UniLoc’s confidence scores can faithfully reflect prediction reliability.

**Figure 3.**
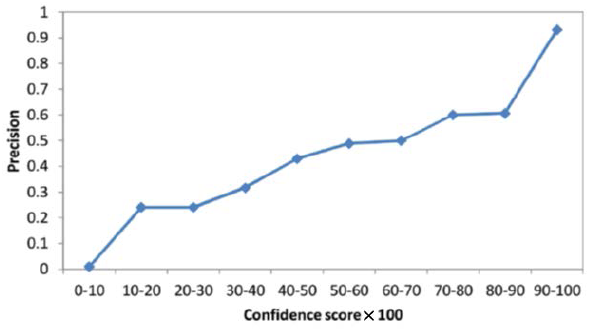
Relationship between confidence scores and precision.

### Match rate and prediction performance

A word related to a target protein contributes to the prediction only if it can be exactly matched to a dictionary entry. Intuitively, the more related words that are matched, the more information UniLoc obtains for the prediction. We define the *match rate* as the ratio of the number of matched entries to the total number of related words of the target protein.Figure 4 shows the relationship between match rate and prediction accuracy on *SP-Euk* and *SP-Prok*. It can be observed that proteins with a higher match rate had better prediction results, suggesting that the prediction performance of UniLoc can be further improved if *RelatedDict* is enriched by adding more annotated proteins and related words.

**Figure 4.**
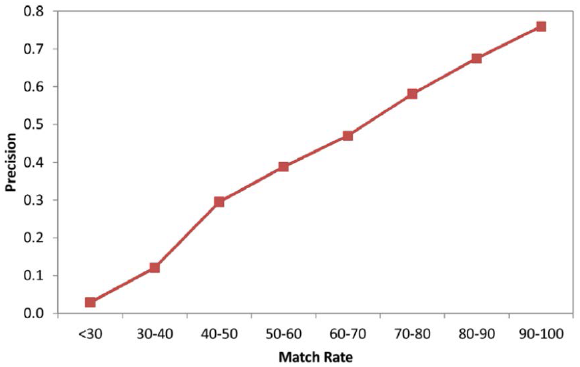
Relationship between match rate and prediction precision. The two factors show a strong positive correlation with a correlation coefficient of 0.99.

### The relationship between intersection percentages and precisions

UniLoc identifies a template protein *q* according to its intersection percentage to the target protein. In general, the higher intersection percentage it has, the more similar it is to the target protein.Figure 5 shows the strong positive relationship between intersection percentage and prediction efficiency (correlation coefficient = 0.93). The average precision is above 60% when the intersection percentage is above 10%. This shows that the intersection percentage can fairly reflect the PSL similarity to the target protein regardless of their sequence identity.

**Figure 5.**
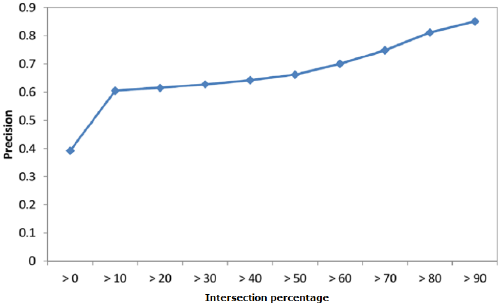
The relationship between intersection percentage and prediction accuracy. The correlation coefficient is 0.93.

### Comparison with existing predictors

We compare the prediction performance of UniLoc with several selected state-of-the-art methods. A PSL predictor was selected if it was built for general eukaryotic or prokaryotic proteins and is available online or as a standalone program. Among the selected methods, some adopt similarity search and some do not. To conduct fair comparisons, all methods are separated by category (eukaryota or prokaryota, denoted as euk/prok) and method (with or without similarity search, denoted as wts/wos). The first group includes YLoc [31] and Euk-mPLoc [28] (euk and wts); the second group includes PSortb [41] (prok and wts); the third includes CELLO[14], Wolf PSORT[18], and SherLoc2 [42] (euk and wos); and the fourth includes CELLO (prok and wos). CELLO provides different prediction modes so it is included in two groups.

Not every selected PSL predictor can handle the whole protein sequences in the benchmark data sets, and some only cover a subset of subcellular locations. Thus each method was only tested on a specialized subset of *SP-Euk*/*SP-Prok* with the locations covered by the method. When UniLoc is compared to PSL predictors with similarity search, a BLAST search for similar proteins is first performed in the SwissProt database. The annotation of the target protein is ignored and its prediction is made based on the top five similar proteins as we do in the PSI-BLAST based prediction method. Table 5, summarizes the comparison between UniLoc and the selected methods in the first and second groups. UniLoc outperforms the selected PSL predictors on both the *SP-Euk* and *SP-Prok* data sets. The average F-scores of UniLoc, Euk-mPLoc2, and YLoc on *SP-Euk* are 84.3, 74.6 and 65.9, respectively; the average F-scores of UniLoc and PSortb on SP-Prok are 88.4 and 79.4, respectively. Table 6, summarizes the comparison between UniLoc and the selected methods in the third and fourth groups. Likewise, UniLoc compares favorably to the selected PSL predictors both on *SP-Euk* and *SP-Prok* for all measures, with the exception that WolfPSORT yields the highest recall as it sacrifices precision. As all selected methods were trained separately based on their own training sets, it is likely that some of the test proteins in the benchmark data sets were already involved in their training process. Thus their evaluation could be overestimated.

**Table 5.** Performance comparison among UniLoc and the selected methods with a homology search step.

| Data set | Predictor | Top1 success rate | Precision | Recall | F-score |
| --- | --- | --- | --- | --- | --- |
| *SP-Euk* | UniLoc | **86.8** | **82.9** | **85.8** | **84.3** |
|  | Euk-mPLoc2 | 80.0 | 79.8 | 70.0 | 74.6 |
|  | YLoc | 65.4 | 63.4 | 68.5 | 65.9 |
| *SP-Prok* | UniLoc | **87.0** | **87.2** | **89.6** | **88.4** |
|  | PSortb | 80.8 | 80.8 | 78.1 | 79.4 |

**Table 6.** Performance comparison among UniLoc and the selected methods which do not use homology search.

| Data set | Predictor | Top1 success rate | Precision | Recall | F-score |
| --- | --- | --- | --- | --- | --- |
| *SP-Euk* | UniLoc | **71.9** | **65.0** | 73.2 | **68.9** |
|  | Wolf PSORT | 64.6 | 49.0 | **81.5** | 61.2 |
|  | CELLO | 63.5 | 62.9 | 56.6 | 59.6 |
|  | SherLoc2 | 64.0 | 64.0 | 51.7 | 57.2 |
| *SP-Prok* | UniLoc | **66.2** | **71.2** | **81.5** | **76.0** |
|  | CELLO | 53.9 | 52.2 | 56.3 | 54.2 |

## CONCLUSIONS

In this study, we demonstrated that UniLoc produced the best performance on the two benchmark data sets compared with several selected PSL methods. We present the concept of related protein words and explore subcellular localization site assignment of an uncharacterized protein by means of identifying its templates. UniLoc identifies template proteins based on the intersection of related words with target proteins.

In summary, we have demonstrated that related words can help to detect the implicit similarity between proteins with the same localization sites, and template proteins with high intersection percentages show a high positive correlation with target proteins. A limitation of UniLoc is the difficulty in producing reliable predictions for proteins which share fewer words with the dictionary. However, this issue will gradually be addressed as more annotated proteins are included each year.

